# Functional dynamics and selectivity of two parallel corticocortical pathways from motor cortex to layer 5 circuits in somatosensory cortex

**DOI:** 10.1101/2024.02.11.579810

**Authors:** Hye-Hyun Kim, Kelly E. Bonekamp, Grant R. Gillie, Dawn M. Autio, Tryton Keller, Shane R. Crandall

**Affiliations:** Department of Physiology, Michigan State University, East Lansing, MI 48824, USA; Molecular, Cellular, and Integrative Physiology Program, Michigan State University East Lansing, MI 48824, USA

## Abstract

In the rodent whisker system, active sensing and sensorimotor integration are mediated in part by the dynamic interactions between the motor cortex (M1) and somatosensory cortex (S1). However, understanding these dynamic interactions requires knowledge about the synapses and how specific neurons respond to their input. Here, we combined optogenetics, retrograde labeling, and electrophysiology to characterize the synaptic connections between M1 and layer 5 (L5) intratelencephalic (IT) and pyramidal tract (PT) neurons in S1 of mice (both sexes). We found that M1 synapses onto IT cells displayed modest short-term depression, whereas synapses onto PT neurons showed robust short-term facilitation. Despite M1 inputs to IT cells depressing, their slower kinetics resulted in summation and a response that increased during short trains. In contrast, summation was minimal in PT neurons due to the fast time course of their M1 responses. The functional consequences of this reduced summation, however, were outweighed by the strong facilitation at these M1 synapses, resulting in larger response amplitudes in PT neurons than IT cells during repetitive stimulation. To understand the impact of facilitating M1 inputs on PT output, we paired trains of inputs with single backpropagating action potentials, finding that repetitive M1 activation increased the probability of bursts in PT cells without impacting the time-dependence of this coupling. Thus, there are two parallel but dynamically distinct systems of M1 synaptic excitation in L5 of S1, each defined by the short-term dynamics of its synapses, the class of postsynaptic neurons, and how the neurons respond to those inputs.

**SIGNIFICANCE STATEMENT:** Normal sensorimotor integration depends in part on the dynamic interactions between the primary motor cortex and the somatosensory cortex, but the functional properties of the excitatory synapses interconnecting the motor cortex with the somatosensory cortex are poorly understood. Our results show that the short-term dynamics of excitatory motor cortex synapses and the nature of the postsynaptic response they generate onto layer 5 pyramidal neurons in the somatosensory cortex depend on the postsynaptic cell type and if their axons project to other cortical areas or subcortical regions. These two parallel but dynamically distinct channels of synaptic excitation constitute previously unknown synaptic circuits by which different temporal patterns of motor cortex activity can shape how signals propagate out of the somatosensory cortex.

## INTRODUCTION

The remarkable ability of the mammalian neocortex to execute complex brain functions depends on long-range interactions between functionally related neocortical regions. Moreover, growing evidence indicates layer 5 (L5) pyramidal neurons, the primary output cells of the neocortex (Wise and Jones, 1977), are a major target of these projections (Petreanu et al., 2009; Dembrow et al., 2015; Kim et al., 2015; Kinnischtzke et al., 2016; Young et al., 2021), and that their activities may play a significant role during defined cortical computations (Xu et al., 2012; Cichon and Gan, 2015; Takahashi et al., 2016; Ranganathan et al., 2018; Takahashi et al., 2020; Mohan et al., 2023). In the rodent whisker sensorimotor system, the whisker primary somatosensory (S1) and motor cortex (M1) are massively interconnected (White and DeAmicis, 1977; Porter and White, 1983; Miyashita et al., 1994; Cauller et al., 1998), and their coordinated activities are critical for normal sensation, motor execution, and sensorimotor integration (Kleinfeld et al., 2006; Diamond et al., 2008).

Neural activity in M1 codes for various whisking parameters (Carvell et al., 1996; Brecht et al., 2004; Hill et al., 2011; Friedman et al., 2012; Huber et al., 2012; Petreanu et al., 2012; Sreenivasan et al., 2016), and the projections from M1 are generally thought to communicate these motor signals to S1, influencing sensorimotor integration and processing in individual neurons (Fee et al., 1997; Petersen, 2019). In S1, the axons of M1 pyramidal neurons arborize in L5/6 and ascend to ramify in L1 (Veinante and Deschenes, 2003). Although previous studies indicate that M1 axons make excitatory connections onto L5 pyramidal neurons in S1 (Petreanu et al., 2009; Rocco-Donovan et al., 2011; Zagha et al., 2013; Kinnischtzke et al., 2014; Kinnischtzke et al., 2016; Martinetti et al., 2022), understanding the nature of M1 influences requires information about the physiological properties of the connections and how L5 cells respond to those signals. A further complication is that L5 pyramidal neurons are grouped into two classes based on their axonal projections (Harris and Shepherd, 2015; Moberg and Takahashi, 2022). Intratelencephalic (IT) neurons project predominately to other neocortical areas, whereas pyramidal tract (PT) neurons project primarily to subcortical structures. In addition, IT and PT neurons have distinct biophysical membrane properties and morphological features that can significantly influence synaptic responses (Wise and Jones, 1977; Magee, 1999; Williams and Stuart, 2000; Hattox and Nelson, 2007; Groh et al., 2010; Morishima et al., 2011; Sheets et al., 2011; Dembrow et al., 2015; Harnett et al., 2015; Kinnischtzke et al., 2016; Anastasiades et al., 2018). They also appear to differ in the efficacy of action potentials (APs) to actively backpropagate into their dendrites (Stuart and Sakmann, 1994; Schiller et al., 1995; Larkum et al., 1999a; Grewe et al., 2010), which may have important implications for how somatically generated APs interact with M1 input (Larkum et al., 1999b). Given these differences and their roles in corticocortical and cortical-subcortical communication, determining the physiological properties of M1 synapses and how L5 neurons respond to those signals is critical for understanding the nature of M1 influences on S1 output.

We previously found that long-range excitatory projections from M1 to S1 generally display synaptic facilitation (Martinetti et al., 2022), a form of short-term plasticity that enhances synaptic transmission when activated repetitively (Jackman and Regehr, 2017). However, we also found that some L5 cells receive depressing M1 synaptic inputs, reducing synaptic strength with activity (Zucker and Regehr, 2002). These differences and the two classes of L5 pyramidal neurons in S1 suggest that M1 synaptic properties could depend on the postsynaptic neuron and its long-range projection target. To test this, we combined optogenetic strategies and retrograde labeling techniques to excite M1 axons within S1, characterize their synaptic properties, and investigate how L5 IT and PT neurons respond to M1 signals. Our results support this hypothesis and suggest that M1 to S1 synaptic circuits are tailored to the divergent functions of the distinct L5 output neurons of S1.

## MATERIALS AND METHODS

### Animals

All experiments were conducted in accordance with the National Institutes of Health (NIH) Guidelines for the Care and Use of Laboratory Animals and approved by the Michigan State University Institutional Animal Care and Use Committee (IACUC). For this study, we used the following mouse lines: CD1 (ICR: Institute for Cancer Research) (Charles River: 022) and the Allen Institute’s Ai14 reporter (Jackson Labs: 007914). Homozygous Ai14 mice were bred with CD1 (ICR) mice, resulting in heterozygous experimental mice for the Cre reporter allele. Animals were group-housed, maintained on a 12:12 hour light-dark cycle, and provided food and water ad libitum. We used both male and female mice in this study.

### Stereotactic Virus Injections

All experiments followed previously described methods for stereotactic injections (Crandall et al., 2017; Martinetti et al., 2022). Briefly, mice (postnatal day 19-60, mean = 26 days) were anesthetized with a Ketaset-Dexdomitor mixture diluted in sterile saline (Ketaset, 70-100 mg/kg; Dexdomitor, 0.25 mg/kg; intraperitoneally). Once deeply anesthetized, eyes were covered with a thin layer of ophthalmic ointment to prevent drying (Patterson Veterinary Artificial Tears), and the mouse was head fixed into a digital stereotaxic setup with a warming base to maintain body temperature (Stoelting). Next, a small craniotomy was made over the injection site, through which viruses or retrograde tracers were pressure injected into the brain via a glass micropipette attached to a Picospritzer pressure system (typically, 0.1-0.2 μL per injection site over 5-10 min). For optogenetic stimulation of M1 axons/terminals, an adeno-associated virus (AAV2) encoding genes for hChR2(H134R)-EYFP fusion proteins (rAAV2/hSyn-hChR2[H134R]-eYFP-WPREpA, AV4384, 3.1-3.5 x 10^12^ viral genomes/ml, University of North Carolina Viral Vector Core) was injected unilaterally into whisker M1. Our previous work defined the center of M1 anatomically to be between 0.9 mm and 1.3 mm anterior and 1.25 mm lateral with respect to bregma (Martinetti et al., 2022), which is consistent with other studies defining whisker M1 in the mouse anatomically and with intracortical microstimulation (Ferezou et al., 2007; Mao et al., 2011; Sreenivasan et al., 2016). For retrograde labeling of PT neurons in S1 projecting to subcortical targets, mice were injected during the same procedure with either cholera toxin subunit B conjugated to Alexa Fluor 647 (CTB647; C34778 Thermo-Fischer) or AAVretro-Cre (pAAV-Ef1a-mCherry-IRES-Cre, 55632-AAVrg, 1.1-3.5 x 10^12^ viral genomes/ml, Addgene) into the ipsilateral posterior medial nucleus of the thalamus (POM), the ipsilateral superior colliculus (SC), or the contralateral spinal trigeminal nucleus (SP5). For retrograde labeling of IT neurons in S1, mice were injected with CTB647 into either M1 (co-injection with the AAV2) or secondary somatosensory cortex (S2). Following injections, the pipette was left in place for an additional 5 min before being slowly withdrawn from the brain, and the scalp closed with a surgical adhesive (GLUture). After surgery, mice were given Antisedan (2.5 mg/kg) to reverse the effects of the Dexdomitor and allowed to recover on a heating pad for 1-2 h before returning to their home cage. Stereotaxic coordinates were relative to bregma: M1 = 1.25 mm lateral, 0.9 and 1.3 mm anterior, and 0.40 and 1.0 mm depth; S2 = 3.6 and 4.2 lateral, 1.1 mm posterior, and 1.95 and 1.16 mm depth; POM = 1.40 mm lateral, 1.27 mm posterior, and 2.84 mm depth; SC = 1.10 mm lateral, 3.23 mm posterior, and 2.25 mm depth; SP5 = 1.8 mm lateral, 6.4 mm posterior, and 4.1 mm depth.

### In vitro slice preparation and live slice imaging

After allowing 21±2 days for ChR2 expression, acute coronal brain slices containing S1 ipsilateral to the M1 injection were prepared for *in vitro* recording and optical stimulation, as previously described (Crandall et al., 2010; Crandall et al., 2015; Martinetti et al., 2022). Briefly, mice (postnatal day 39-81, mean = 46 days) were deeply anesthetized by inhalation of isoflurane before being decapitated. After decapitation, the brain was immediately removed and submerged in an ice-cold (∼4°C) oxygenated (95% O_2_, 5% CO_2_) cutting solution containing the following (in mM): 3 KCl, 1.25 NaH_2_PO_4_, 10 MgSO_4_, 0.5 CaCl_2_, 26 NaHCO_3_, 10 glucose and 234 sucrose. Brain slices (300 μm thick) were prepared using a vibrating tissue slicer (VT1200S, Leica Microsystems) and then transferred to an incubation chamber filled with warm (32°C) oxygenated artificial cerebrospinal fluid (ACSF) containing (in mM): 126 NaCl, 3 KCl, 1.25 NaH_2_PO_4_, 2 MgSO_4_, 2 CaCl_2_, 26 NaHCO_3_, and 10 glucose. Slices were allowed to recover at 32°C for 20 min and then at room temperature for a minimum of 40 min. Before recording, all live brain slices were imaged using a Zeiss Axio Examiner.A1 microscope equipped with a 2.5x objective (Zeiss EC plan-Neofluar) and an Olympus XM101R camera with cellSens software. Bright-field and fluorescence images were taken to confirm the accuracy of the injections, tissue health, and the overall expression of M1 axons/terminals and retrogradely filled cells in S1. We only recorded slices obtained from mice without signs of damage to the brain or off-target injections. *In vitro electrophysiological recordings and data acquisition.* Brain slices were transferred to a submersion-type recording chamber continuously perfused (2-3 mL/min) with warm (32±1 °C) oxygenated ACSF containing (in mM): 126 NaCl, 3 KCl, 1.25 NaH_2_PO_4_, 1 MgSO_4_, 1.2 CaCl_2_, 26 NaHCO_3_ and 10 glucose. Neurons were visualized using infrared differential interference contrast optics (IR-DIC) and fluorescence microscopy using a Zeiss Axio Examiner.A1 microscope equipped with a 40x water immersion objective (Zeiss, W plan-Apo 40x/1.0 NA) and video camera (Olympus, XM10-IR). All electrophysiological recordings were made from the soma using borosilicate glass patch pipettes (tip resistance 3-6 MΩ) filled with a potassium-based internal solution containing (in mM): 130 K-gluconate, 4 KCl, 2 NaCl, 10 HEPES, 0.2 EGTA, 4 ATP-Mg, 0.3 GTP-Tris and 14 phosphocreatine-K (pH 7.25, 290 mOsm). In some experiments, neurobiotin (0.25%, w/v; Vector Laboratories) was added to the internal solution to fill individual cells for subsequent morphological reconstruction. Cells were filled for 20-60 min before placing the slice in 4% paraformaldehyde in 0.1 M phosphate buffer solution. All voltages were corrected for a -14 mV liquid junction potential.

Electrophysiological data were acquired and digitized at 20 kHz using Molecular Devices hardware and software (MultiClamp 700B amplifier, Digidata 1550B4, pClamp 11). Signals were low-pass filtered at 10 kHz (current-clamp) or 3 kHz (voltage-clamp) before digitizing. During whole-cell recordings, the pipette capacitances were neutralized, and series resistances were compensated online (100% for current-clamp and 60-80% for voltage-clamp). Series resistances were continually monitored and adjusted during experiments to ensure sufficient compensation. ZD7288 (Tocris Cat. No. 1000) was prepared as recommended by the manufacturer and diluted in ACSF just before use. To minimize the off-target effects of the agent, a low concentration of ZD7288 (10 μM) was bath applied for a maximum period of 5 minutes (Harnett et al., 2015; Hsu et al., 2018) before subsequent experimental tests.

### Photostimulation

ChR2 was optically excited using a high-power white light-emitting diode (LED) (Mightex LCS-5500-03-22) driven by a Mightex LED controller (SLC-AA02-US). Collimated light was reflected through a single-edge dichroic beam-splitter (Semrock FF660-FDi02) and then a high magnification water immersion objective (Zeiss, W-Plan-Apo 40x/1.0 NA), resulting in an estimated spot diameter of ∼1500 um and maximum LED power at the focal plane of ∼32 mW (∼18 mW/mm^2^). Optical stimuli were delivered as 0.5 ms flashes and were directed at ChR2-expressing axons/terminals by centering the light over the cell bodies. LED on/off times were fast (< 50 μs) and of constant amplitude and duration when verified with a fast photodiode (Thorlabs, DET36A). The LED intensity was typically adjusted to evoke 100-200 pA EPSC in the recorded neuron when held in voltage-clamp at -94 mV, near the reversal potential for inhibition (Martinetti et al., 2022).

*Histology, imaging, and neuronal reconstruction.* Tissue for histology was prepared from acute coronal brain slices, as previously described (Crandall et al., 2017; Martinetti et al., 2022). Briefly, 300 μm thick coronal brain slices containing S1 were transferred to a 4% paraformaldehyde in 0.1 M phosphate buffer solution overnight at 4°C (18–24 hr). The next day, slices were changed to a 30% sucrose in 0.1 M phosphate buffer solution until re-sectioned (4°C; 2–3 days). Individual brain slices were re-sectioned at 50-100 µm using a freezing microtome (Leica SM2010 R). Sections were washed 2 times in 0.1 M phosphate buffer followed by 3 washes in 0.1 M phosphate buffer with 0.15 M NaCl, pH 7.4 (PBS, 5 min/wash). After washing, sections were incubated for 1 hr at room temperature in a blocking solution containing 0.1% Tween, 0.25% Triton X-100, 10% normal goat serum in PBS. Sections were then incubated using Streptavidin/Biotin Blocking Kit (Vector Labs, Cat # SP-2002), 30 minutes in streptavidin solution, followed by 30 minutes in biotin solution with a brief rinse of PBS after each. Sections were then incubated with Alexa Fluor 647-conjugated streptavidin (Thermo-Fisher Scientific, Cat #S21374, 1:1000, 2mg/ml) solution prepared in blocking solution for 3 hrs with rotation at room temperature. Following incubation, sections were washed 3 times in PBS and 2 times in 0.1 M phosphate buffer solution (10 min/wash) before being mounted and cover-slipped using Vectashield Vibrance w/DAPI (Vector Laboratories H-1800). Fluorescent conjugates of streptavidin were stored and prepared as recommended by the manufacturer. Neurobiotin-filled cells were initially identified and assessed using a Zeiss Axio Imager.D2 fluorescence microscope. If suitable, confocal image stacks of labeled neurons were taken using a Nikon A1 Laser Scanning Confocal Microscope with a 20X Plan apochromat Lambda D NA 0.8 objective and laser excitation 405nm, 488nm, 561nm, and 647nm. Image brightness and contrast were adjusted offline using Fiji software (Schindelin et al., 2012). All filled cells were reconstructed in three dimensions using the Simple Neurite Tracer Plugin within FIJI software. The tuft dendrites of reconstructed neurons were analyzed using the Path Manager’s measurement feature in Simple Neurite Tracer, with the total path length (cable length) defined as the sum of the distances along each tuft dendritic segment within L1.

*Data analysis.* Analysis of electrophysiological data was performed in Molecular Devices Clampfit 11, Microsoft Excel, and Igor Pro. The synaptic responses to optical stimulation were recorded from postsynaptic neurons in whole-cell voltage-clamp and current-clamp, and the amplitude of an evoked EPSC/EPSP was measured relative to a baseline (1-10 ms) before the stimulus. The maximum depolarization of an evoked EPSP was measured relative to the pre-train baseline (10 ms). The amplitude was typically measured over the 10 ms immediately after stimulus onset, but in some cases, a 4.5 to 5 ms window was used. Intrinsic physiological properties were measured from rest using previously described methods (Crandall et al., 2017). Bursts were defined as 3 or more action potentials riding upon a slow depolarizing envelope with a mean inter-spike frequency greater than 150 Hz (Connors et al., 1982; McCormick et al., 1985). Cell depth measurements were calculated as the normalized distance from pia to white matter across the cortical column.

*Statistical analysis.* Statistical comparisons were performed in OriginPro 2019. The Shaprio-Wilk test was first applied to determine whether the data had been drawn from a normally distributed population; in this case, parametric tests were used (Paired t-test or two-sample t-test). If the assumption of normality was invalid, nonparametric tests were used (Wilcoxon paired signed-rank test or Mann-Whitney U test). A one-way analysis of variance (ANOVA) was used for multiple comparisons, and a two-way ANOVA was used to compare short-term synaptic dynamics across 20 Hz trains. All tests were two-tailed. All the data are presented as mean ± SEM, and statistical significance was defined as p < 0.05. No sex differences were found in the initial EPSC amplitude (when normalized to L2/3 excitatory cells), paired-pulse ratio, or EPSC peak ratio for the tenth response in a 20 Hz train (stim10/ stim1) for either the IT (M1p and S2p) or PT (SCp, SP5p, and POMp) populations, so the data from males and females were combined (p > 0.15079, Mann-Whitney U Test or two-sample t-test; data not shown).

## RESULTS

To investigate M1 synaptic responses in L5 pyramidal neurons of S1 with distinct cortical or subcortical targets, we injected fluorescent retrograde tracers in combination with an adeno-associated virus (AAV2) carrying genes for a Channelrhodospin-2/enhanced yellow fluorescent protein (ChR2/EYFP) into M1 of Ai14 reporter mice *in vivo*. For most experiments, a far-red cholera toxin subunit B (CTB647) was injected into the ipsilateral M1 or secondary somatosensory cortex (S2) to label IT neurons or an AAVretro-Cre (Tervo et al., 2016) was injected into the ipsilateral posterior medial nucleus of the thalamus (POM), ipsilateral superior colliculus (SC), or contralateral spinal trigeminal nucleus (SP5) to label PT neurons. We chose CTB647 to label IT neurons because we observed minimal labeling in L5 using AAVretro-Cre (n = 5 mice; **Extended Data** Figure 1). Three weeks after injections, retrogradely labeled cells and ChR2/EYFP-expressing M1 axons/terminals were visible within S1 (**Figure 1A**). M1 terminal arbors were densest in L1 and L5/6 of S1, consistent with previous studies (Veinante and Deschenes, 2003; Martinetti et al., 2022). Examining the labeled neurons revealed significant electrophysiological, anatomical, and morphological differences consistent with those previously reported for identified L5 IT and PT populations (Wise and Jones, 1977; Chagnac-Amitai et al., 1990; Mason and Larkman, 1990; Kasper et al., 1994; Hattox and Nelson, 2007; Groh et al., 2010; Morishima et al., 2011; Oberlaender et al., 2012; Kinnischtzke et al., 2016). Specifically, the input resistance of PT cells was smaller, input capacitance was larger, and membrane time constant was shorter when compared to IT cells (**Figure 1B**, **Table 1**). PT neurons had a more depolarized resting membrane potential, lower spike threshold, lower rheobase, and narrower spikes than IT cells. In addition, while most IT neurons could be classified as regular spiking (RS) neurons (96%), exhibiting robust spike frequency adaptation, many PT neurons (40.5%) could be characterized as intrinsically bursting (Connors et al., 1982; McCormick et al., 1985), responding to positive current injections with high-frequency bursts of action potentials. Anatomically, PT neurons were also located primarily in the lower part of L5 (L5b). Lastly, while both cell types had L1 projecting apical dendrites, PT neurons had large diameter apical dendrites with extensive branching in L1 (i.e., thick-tufted), whereas IT neurons had smaller diameter apical dendrites and less branching in L1 (i.e., thin-tufted) (Total path length of L1 tufts; M1p: 963.6 ± 167.4 μm, n = 3; S2p: 681.8 ± 74.0 μm, n = 2; SCp: 1256.8 μm, n = 1; SP5p: 2327.9 μm, n = 1; POMp: 2323.7 ± 456.5 μm, n = 7; **Figure 1C**). These anatomical and physiological results indicate that our cells were not unhealthy or damaged due to the retrograde labeling strategy or the slicing procedure.

**Figure 1.**
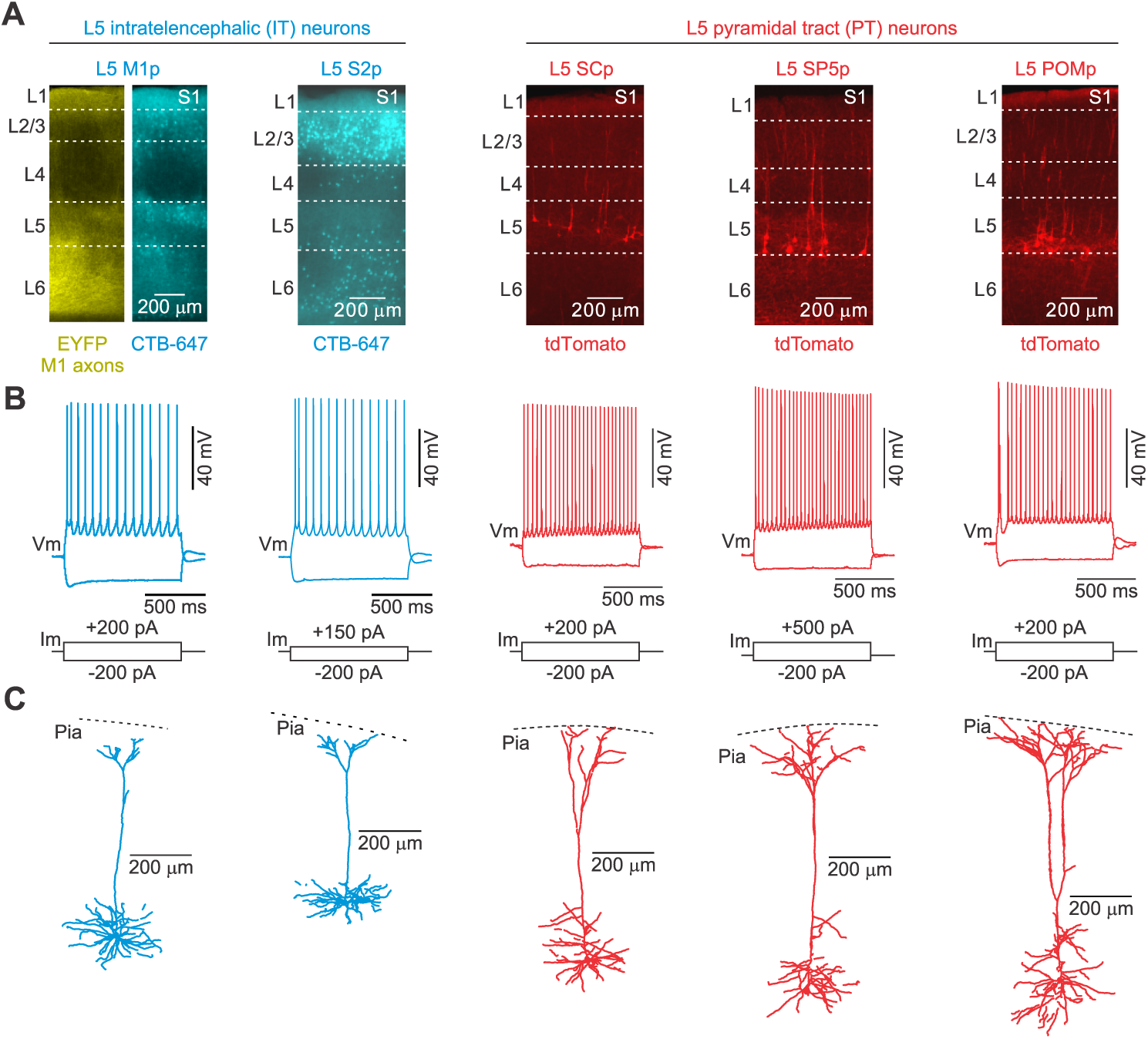
ChR2-EYFP expression in M1 projection neurons and retrograde identification of distinct L5 projection neurons in the mouse S1. **A**, AAV-encoding ChR2-EYFP was injected into the ipsilateral M1 to label their axonal projections in S1. Retrograde fluorescent tracers (CTB-647 or rAAV-Cre) were also injected into M1 or S2 to label subclasses of IT neurons, or SC, SP5, or POM to label different subclasses of PT neurons. EYFP expression in M1 axons/terminals (far left image) and retrogradely labeled neurons in L5 were evident 3 weeks post-injection. **B-C**, Whole-cell recordings (**B**) and neurobiotin filling (**C**) of L5 pyramidal neurons not only revealed that labeled cells were healthy and had apical dendrites that terminated in tufts near the pia but also confirmed the physiological and morphological differences between L5 projection classes. See **Table 1** for physiological differences.

**Table 1.**
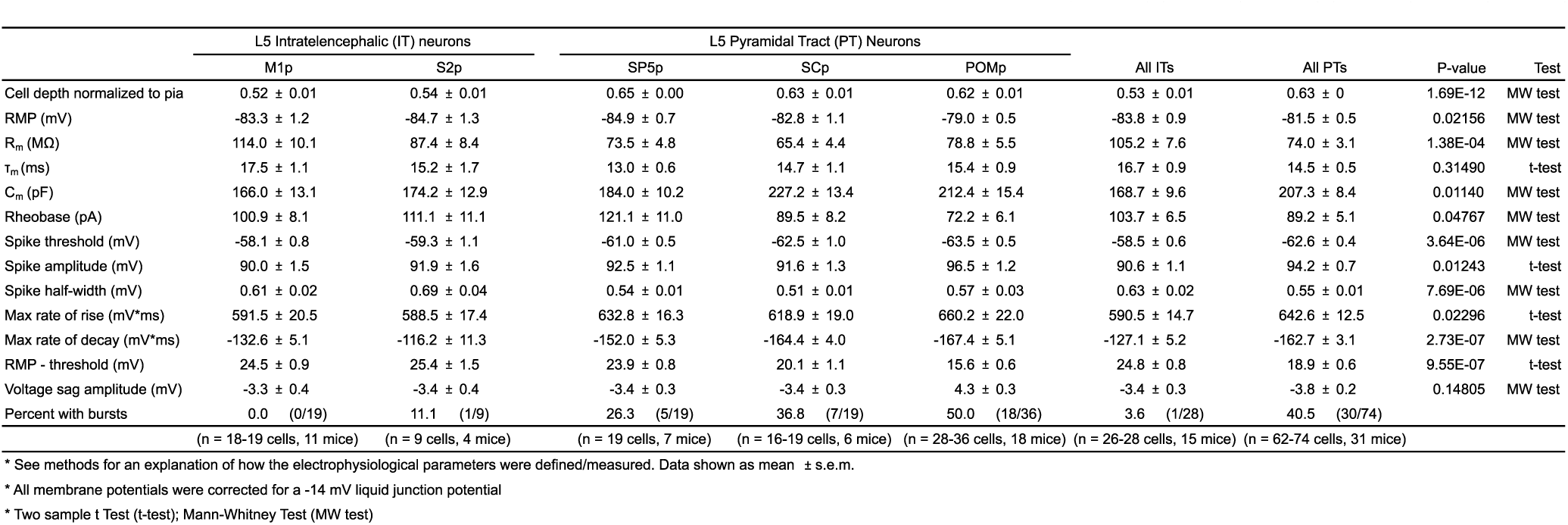
Electrophysiological properties of excitatory projection neurons in LS of mouse primary somatosensory cortex.

|  | L5 Intratelencephalic (IT) neurons |  | L5 Pyramidal Tract (PT) Neurons |  |  | All ITs | All PTs | P-value | Test |
| --- | --- | --- | --- | --- | --- | --- | --- | --- | --- |
|  | M1p | S2p | SP5p | SCp | POMp |  |  |  |  |
| Cell depth normalized to pia | 0.52 ± 0.01 | 0.54 ± 0.01 | 0.65 ± 0.00 | 0.63 ± 0.01 | 0.62 ± 0.01 | 0.53 ± 0.01 | 0.63 ± 0 | 1.69E-12 | MW test |
| RMP (mV) | -83.3 ± 1.2 | -84.7 ± 1.3 | -84.9 ± 0.7 | -82.8 ± 1.1 | -79.0 ± 0.5 | -83.8 ± 0.9 | -81.5 ± 0.5 | 0.02156 | MW test |
| R<sub>m</sub> (MΩ) | 114.0 ± 10.1 | 87.4 ± 8.4 | 73.5 ± 4.8 | 65.4 ± 4.4 | 78.8 ± 5.5 | 105.2 ± 7.6 | 74.0 ± 3.1 | 1.38E-04 | MW test |
| τ<sub>m</sub> (ms) | 17.5 ± 1.1 | 15.2 ± 1.7 | 13.0 ± 0.6 | 14.7 ± 1.1 | 15.4 ± 0.9 | 16.7 ± 0.9 | 14.5 ± 0.5 | 0.31490 | t-test |
| C<sub>m</sub> (pF) | 166.0 ± 13.1 | 174.2 ± 12.9 | 184.0 ± 10.2 | 227.2 ± 13.4 | 212.4 ± 15.4 | 168.7 ± 9.6 | 207.3 ± 8.4 | 0.01140 | MW test |
| Rheobase (pA) | 100.9 ± 8.1 | 111.1 ± 11.1 | 121.1 ± 11.0 | 89.5 ± 8.2 | 72.2 ± 6.1 | 103.7 ± 6.5 | 89.2 ± 5.1 | 0.04767 | MW test |
| Spike threshold (mV) | -58.1 ± 0.8 | -59.3 ± 1.1 | -61.0 ± 0.5 | -62.5 ± 1.0 | -63.5 ± 0.5 | -58.5 ± 0.6 | -62.6 ± 0.4 | 3.64E-06 | MW test |
| Spike amplitude (mV) | 90.0 ± 1.5 | 91.9 ± 1.6 | 92.5 ± 1.1 | 91.6 ± 1.3 | 96.5 ± 1.2 | 90.6 ± 1.1 | 94.2 ± 0.7 | 0.01243 | t-test |
| Spike half-width (mV) | 0.61 ± 0.02 | 0.69 ± 0.04 | 0.54 ± 0.01 | 0.51 ± 0.01 | 0.57 ± 0.03 | 0.63 ± 0.02 | 0.55 ± 0.01 | 7.69E-06 | MW test |
| Max rate of rise (mV/ms) | 591.5 ± 20.5 | 588.5 ± 17.4 | 632.8 ± 16.3 | 618.9 ± 19.0 | 660.2 ± 22.0 | 590.5 ± 14.7 | 642.6 ± 12.5 | 0.02296 | t-test |
| Max rate of decay (mV/ms) | -132.6 ± 5.1 | -116.2 ± 11.3 | -152.0 ± 5.3 | -164.4 ± 4.0 | -167.4 ± 5.1 | -127.1 ± 5.2 | -162.7 ± 3.1 | 2.73E-07 | MW test |
| RMP - threshold (mV) | 24.5 ± 0.9 | 25.4 ± 1.5 | 23.9 ± 0.8 | 20.1 ± 1.1 | 15.6 ± 0.6 | 24.8 ± 0.8 | 18.9 ± 0.6 | 9.55E-07 | t-test |
| Voltage sag amplitude (mV) | -3.3 ± 0.4 | -3.4 ± 0.4 | -3.4 ± 0.3 | -3.4 ± 0.3 | 4.3 ± 0.3 | -3.4 ± 0.3 | -3.8 ± 0.2 | 0.14805 | MW test |
| Percent with bursts | 0.0 (0/19) | 11.1 (1/9) | 26.3 (5/19) | 36.8 (7/19) | 50.0 (18/36) | 3.6 (1/28) | 40.5 (30/74) |  |  |
|  | (n = 18-19 cells, 11 mice) | (n = 9 cells, 4 mice) | (n = 19 cells, 7 mice) | (n = 16-19 cells, 6 mice) | (n = 28-36 cells, 18 mice) | (n = 26-28 cells, 15 mice) | (n = 62-74 cells, 31 mice) |  |  |

\* See methods for an explanation of how the electrophysiological parameters were defined/measured. Data shown as mean ± s.e.m.
\* All membrane potentials were corrected for a -14 mV liquid junction potential
\* Two sample t Test (t-test); Mann-Whitney Test (MW test)

To compare the synaptic properties of M1 inputs onto L5 IT and PT neurons in S1, we first used whole-cell voltage clamp recordings to measure monosynaptic excitatory currents evoked by full-field photostimulation of M1 axons/terminals while holding the membrane potential near the reversal for inhibition (-94 mV). To control for variability in ChR2 expression across slices and mice, we compared synaptic dynamics of the M1 synapses onto each L5 neuron to those contacting L2/3 RS neuron because our previous work has shown that light stimulation of M1 arbors evokes facilitating synaptic excitation in these neurons (Martinetti et al., 2022). Pairs were recorded sequentially and in varying order from cells in the same slice and column. When using 20 Hz optical stimuli (paired-pulse and trains), we found stimulation drove short-latency (∼2 ms) monosynaptic excitatory postsynaptic currents (EPSCs) with short-term synaptic dynamics that depended on the class of L5 neuron. EPSCs recorded in M1 and S2 projecting (M1p and S2p, respectively) neurons remained relatively unchanged or depressed weakly, while SCp, SP5p, and POMp responses exhibited synaptic facilitation (**Figure 2**). Although short-term plasticity was significantly different for both M1p and S2p neurons compared to control L2/3 RS cells, M1 responses in all 3 classes of PT neurons facilitated similarly to L2/3 neurons, increasing 57-86% across trains. Within projection classes, we observed no significant difference between IT cells (M1p and S2p) or PT neurons (SCp, SP5p, and POMp) in initial EPSC amplitude (when normalized to L2/3 excitatory cells), paired-pulse ratio, or EPSC peak ratio for the tenth response in a 20 Hz train (stim_10_/ stim_1_; **Figure 2 K, L**). Collapsing the data from multiple subclasses into IT and PT groups revealed that despite equivalent synaptic strength, when normalized to their paired L2/3 RS cell, the short-term synaptic dynamics of the M1-evoked EPSCs were significantly different between L5 IT and PT neurons (**Figure 3A, B**).

**Figure 2.**
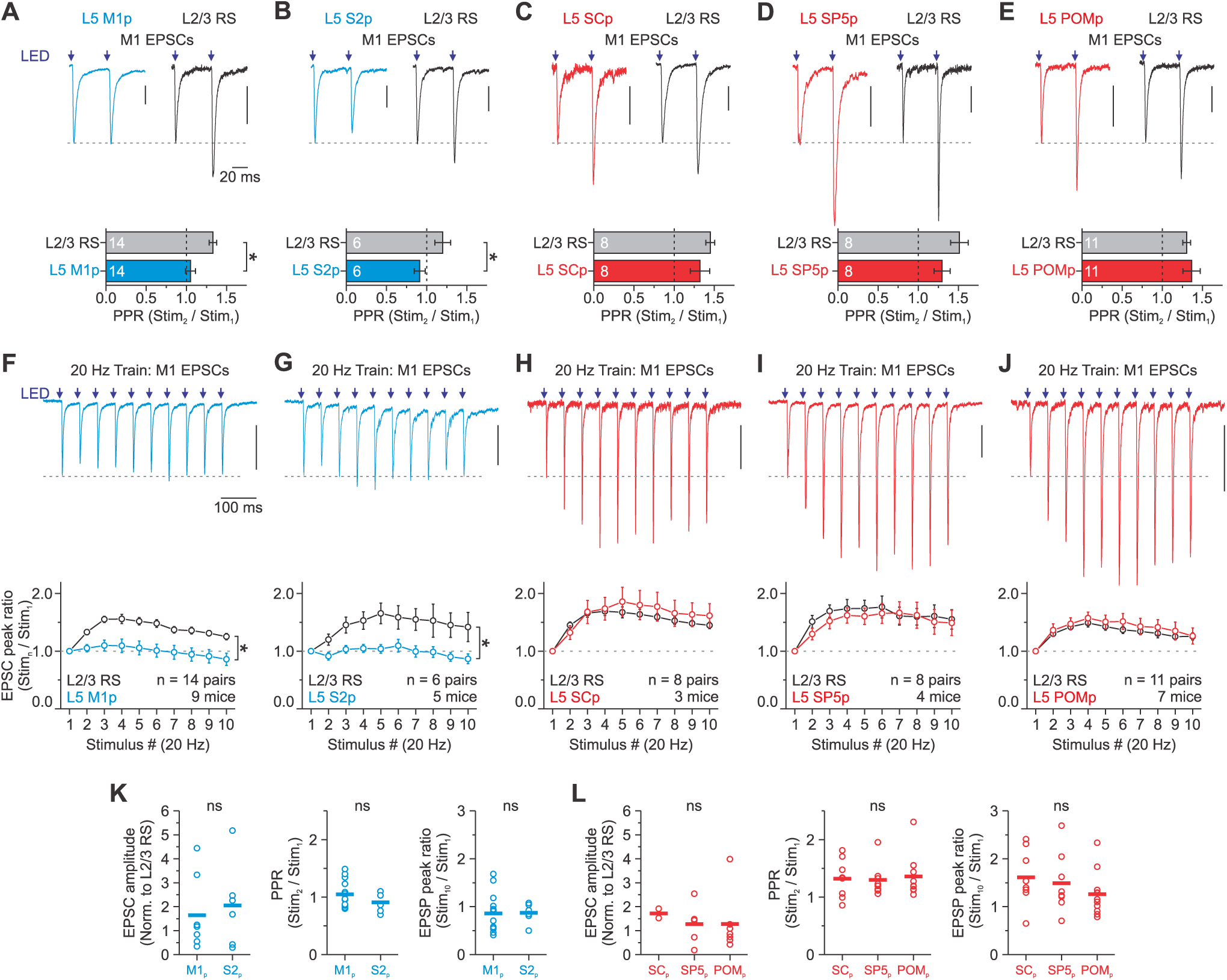
L5 projection neurons in S1 are excited differently by long-range M1 inputs. **A-E**, Top: Average EPSCs evoked by two optical stimuli (LED, Purple arrows, 0.5 ms pulse duration) delivered at a 50 ms interval (20 Hz). Data are shown from L2/3 neurons paired to various L5 IT and PT subtypes, including L5 M1p (**A**), L5 S2p (**B**), L5 SCp (**C**), L5 SP5p (**D**), L5 POMp (**E**). Vertical scale bars, 50 pA (**A-E**). Bottom: Summary of paired-pulse ratios for a 50-ms interstimulus interval. Asterisks denote a significant difference from responses evoked in control L2/3 RS neurons (M1p: 1.05 ± 0.06, L2/3: 1.33 ± 0.05, n = 14 pairs, 9 mice; p = 0.0073, paired t-test; S2p: 0.91 ± 0.07, L2/3: 1.20 ± 0.10, n = 6 pairs, 5 mice; p = 0.00329, paired t-test; SCp: 1.32 ± 0.12, L2/3: 1.45 ± 0.06, n = 8 pairs, 4 mice; p = 0.34802, paired t-test; SP5p: 1.30 ± 0.10, L2/3: 1.51 ± 0.11, n = 8 pairs, 4 mice; p = 0.18343, Wilcoxon paired signed-rank test; POMp: 1.36 ± 0.05, L2/3: 1.30 ± 0.0.11, n = 11 pairs, 7 mice; p = 1.0, Wilcoxon paired signed-rank test). **F-J**, Top: Average EPSCs evoked by a 20 Hz train of optical stimuli recorded in a single L5 M1p neuron (**F**), a L5 S2p neuron (**G**), a L5 SCp neuron (**H**), a L5 SP5p neuron (**I**), and a L5 POMp neuron (**J**). Vertical scale bars, 100 pA (**F-J**). Bottom: Summary of EPSC amplitudes plotted as a function of stimulus number within 20 Hz trains for all L2/3–L5 pairs (normalized to first responses) (M1p vs. L2/3: p = 2.88E-20; S2p vs. L2/3: p = 4.37E-10; SCp vs. L2/3: p = 0.18018; SP5p vs. L2/3: p = 0.24162; POMp vs. L2/3: p = 0.09125; two-way ANOVA, stim. 2–10). EPSCs were recorded at -94 mV in voltage-clamp, near the reversal for inhibition, and the light intensity for each cell was set to obtain an initial peak of 100-200 pA. **K**, Comparison of initial EPSC amplitude (normalized to L2/3 response), paired-pulse ratio, and EPSC ratio for the tenth pulse in a 20 Hz train for the IT subclasses M1p and SC2 (EPSP amplitude, p = 0.74689, Mann-Whitney U test; PPR, p = 0.20283, two-sample t-test; stim10/stim1, p = 0.96999, two-sample t-test). **L**, Same as (**K**) but for the PT subclasses SCp, SP5p, and POMp (EPSP amplitude, p = 0.0.86112, one-way ANOVA; PPR, p = 0.90595, one-way ANOVA; stim10/stim1, p = 0.38276, one-way ANOVA). Values are represented as mean ± SEM.

**Figure 3.**
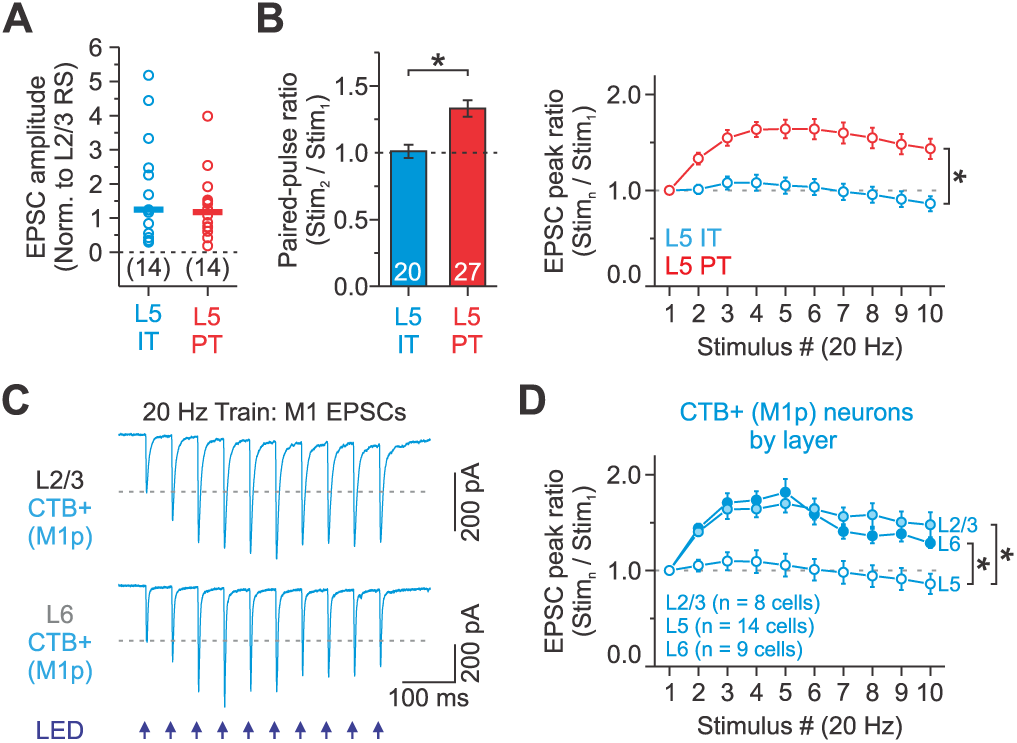
Synaptic responses during repetitive M1 activation are less facilitating for L5 IT than L5 PT neurons. **A**, L5 IT and PT neurons receive similar strength M1 inputs. EPSCs of both cells were normalized to control L2/3 RS neurons (IT: 14 cells, 11 mice; PT: 14 cells, 10 mice; p = 0.66247, Mann-Whitney U test). Bars represent the mean (**A**). We normalized the evoked response in a given L5 cell type to the response in the L2/3 neuron to control for the variability in the level of ChR2 expression in different animals. **B**, Summary of short-term plasticity for all L5 IT (M1p and S2p) and PT neurons (SCp, SP5p, and POMp) to a pair of 20 Hz optical stimuli (left) and a 20 Hz optical stimulus train (right) (data combined from Figure 2) (PPR: L5 IT: 1.01 ± 0.05, n = 20 cells, 15 mice; L5 PT: 1.33 ± 0.06, n = 27 cells, 15 mice; p = 1.89E-4, Mann-Whitney U Test; Train: p = 1.53E-30, two-way ANOVA, stim. 2–10). **C**, Average EPSCs evoked by a 20 Hz train of optical stimuli recorded in a single L2/3 and L6 M1p neuron labeled with CTB. **D**, Summary of EPSC amplitudes plotted as a function of stimulus number within 20 Hz trains for CTB-labeled M1p neurons in L2/3 (n = 8 cells, 3 mice), L5 (n = 14 cells, 9 mice), and L6 (n = 9 cells, 4 mice) (normalized to first responses) (* indicates p < 2.23E-18, two-way ANOVA, stim. 2–10). L5 M1p data from Figure 2. Values are represented as mean ± SEM (**B, D**).

To rule out the possibility that CTB caused abnormal short-term dynamics, we compared M1-evoked EPSCs in CTB-labeled M1p cells across multiple layers in S1. In contrast to M1p cells in L5, the EPSCs of M1p cells in L2/3 and L6 showed clear short-term facilitation (**Figure 3C, 3D**). Moreover, the magnitude of the short-term facilitation was similar to that observed in L5 PT neurons, increasing 70-82% across trains (**Figure 3B, 3D**). Therefore, CTB does not appear to cause abnormal short-term dynamics in L5 IT neurons.

These data confirm that long-range synapses from M1 to L5 of S1 excite both IT and PT neurons but reveal target-cell-specific differences in short-term plasticity. Specifically, the M1 to IT synapse displays modest frequency-dependent depression, whereas the M1 to PT synapse shows robust frequency-dependent facilitation.

To determine the functional impact of differences in synaptic properties and intrinsic physiology, we next examined M1 synaptic responses in current-clamp. Optical stimuli were again delivered full-field to active ChR2-expressing M1 arbors in S1 while controlling for the amplitude of the synaptic potential (∼3 mV). Furthermore, all cells were injected with a negative current to hold their membrane potential near the reversal for inhibition, minimizing any impact of di-synaptic inhibition without adding GABA_A_ and GABA_B_ receptor blockers to the external solution. Note that all L5 IT data was collected from only M1p cells from this point forward. When recording from the soma, we found that the time course of M1-evoked responses for PT and IT neurons were different, with excitatory postsynaptic potentials (EPSPs) in PT neurons having significantly faster rise times (20-80%), shorter half-widths, and faster decays (**Figure 4A**). We observed no difference in the time course of EPSPs across PT subclasses (p > 0.45173, one-way ANOVA). Previous studies have shown that hyperpolarization-activated cation (HCN) channels underlying h-current (Ih) can significantly impact the time course of subthreshold PSPs in L5 PT neurons due to their increased Ih expression (Magee, 1999; Williams and Stuart, 2000; Sheets et al., 2011; Dembrow et al., 2015; Harnett et al., 2015; Anastasiades et al., 2018). Indeed, in the presence of ZD-7288 (10 μM) to block HCN channels, the change in M1-evoked EPSP decay tau was not significant in IT neurons but robust in PT neurons (**Figure 4B)**. Together, these data indicate that the time course of M1-evoked EPSPs is faster in L5 PT than IT neurons and that the faster kinetics are at least partially attributable to differences in their Ih, consistent with reports examining other synaptic inputs to L5 neurons (Sheets et al., 2011; Dembrow et al., 2015; Anastasiades et al., 2018).

**Figure 4.**
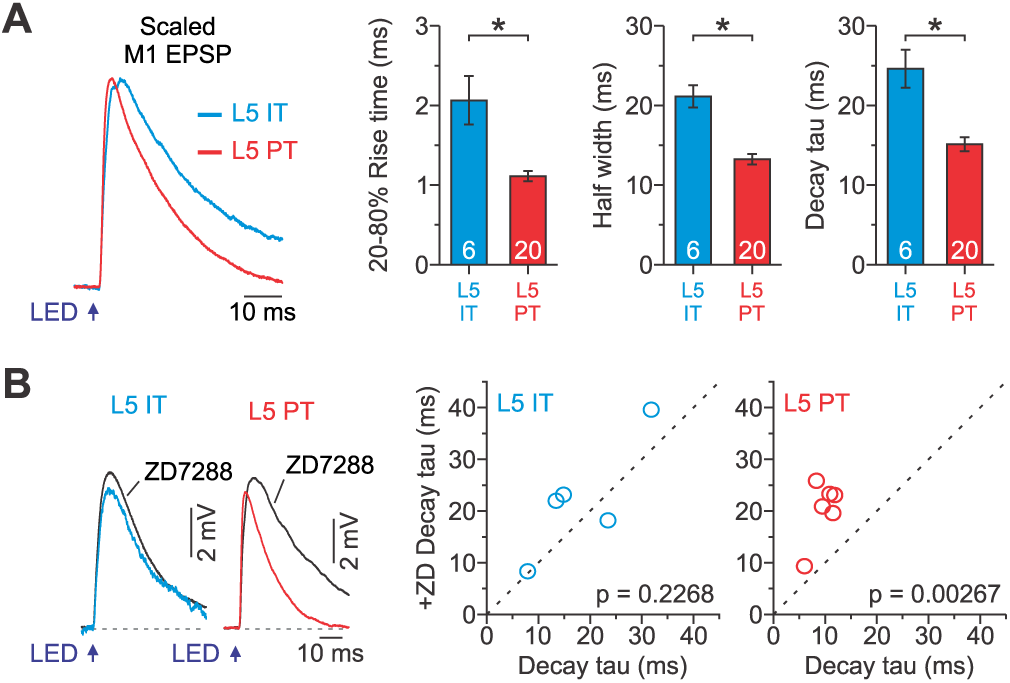
Time course of M1 synaptic responses at L5 PT and IT neurons. **A**, Left: Average M1-evoked EPSP, scaled to match amplitude, recorded in L5 IT and PT neurons. Right: Summary plots showing the kinetics of M1-evoked EPSPs for both cell types, as measured by the 20-80% rise time (IT: 2.1 ± 0.3 ms; PT: 1.1 ± 0.6 ms; p = 0.00113, Mann-Whitney U test), half-width (IT: 21.1 ± 1.4 ms; PT: 13.2 ± 0.7 ms; p = 1.02E-5, two-sample t-test), and decay tau (IT: 24.6 ± 2.4 ms; PT: 15.1 ± 0.9 ms; p = 1.0E-4, two-sample t-test, n = 6 IT cells from 3 mice; n = 20 PT cells from 11 mice). **B,** Left: Average M1-evoked EPSP recorded under control conditions and in the presence of ZD7288 (10 μM) for L5 IT and PT neurons. Right: Summary plots showing the change in decay tau for both cell types (IT control: 18.3 ± 4.2 ms, IT +ZD: 22.2 ± 5.01 ms, n = 5 cells from 3 mice, p = 0.2268, Paired t-test; PT control: 9.7 ± 0.9 ms, PT +ZD: 20.4 ± 2.4 ms, n= 6 cells from 3 mice, p = 0.00267, Paired t-test). Values are represented as mean ± SEM.

Next, we wanted to investigate how L5 neurons responded during repetitive M1 activation. When we delivered optical stimuli in 20-Hz trains, we found that M1 stimulation evoked individual EPSPs in IT neurons that depressed weakly, whereas the EPSPs in PT neurons facilitated (EPSP peak measured from a 1 ms baseline before each light pulse; **Figure 5A-D**). This pattern is highly consistent with the synaptic currents described above. There was no difference in the magnitude of facilitation across PT subclasses (Stimulus 10/ Stimulus 1; p = 0.38114, one-way ANOVA) or the EPSP time course within trains (Decay tau; p < 0.20318, paired t-test). However, because the time course of the elicited EPSPs was slower in IT neurons, we observed a significant overlap of consecutive EPSPs throughout the 20-Hz train, resulting in a peak depolarization that increased during the train when compared to the individual EPSPs peaks (**Figure 5C**). The peak depolarization was measured from a 10 ms pre-train baseline, which accounts for the sum of the response from prior pulses plus the latest EPSP.

**Figure 5.**
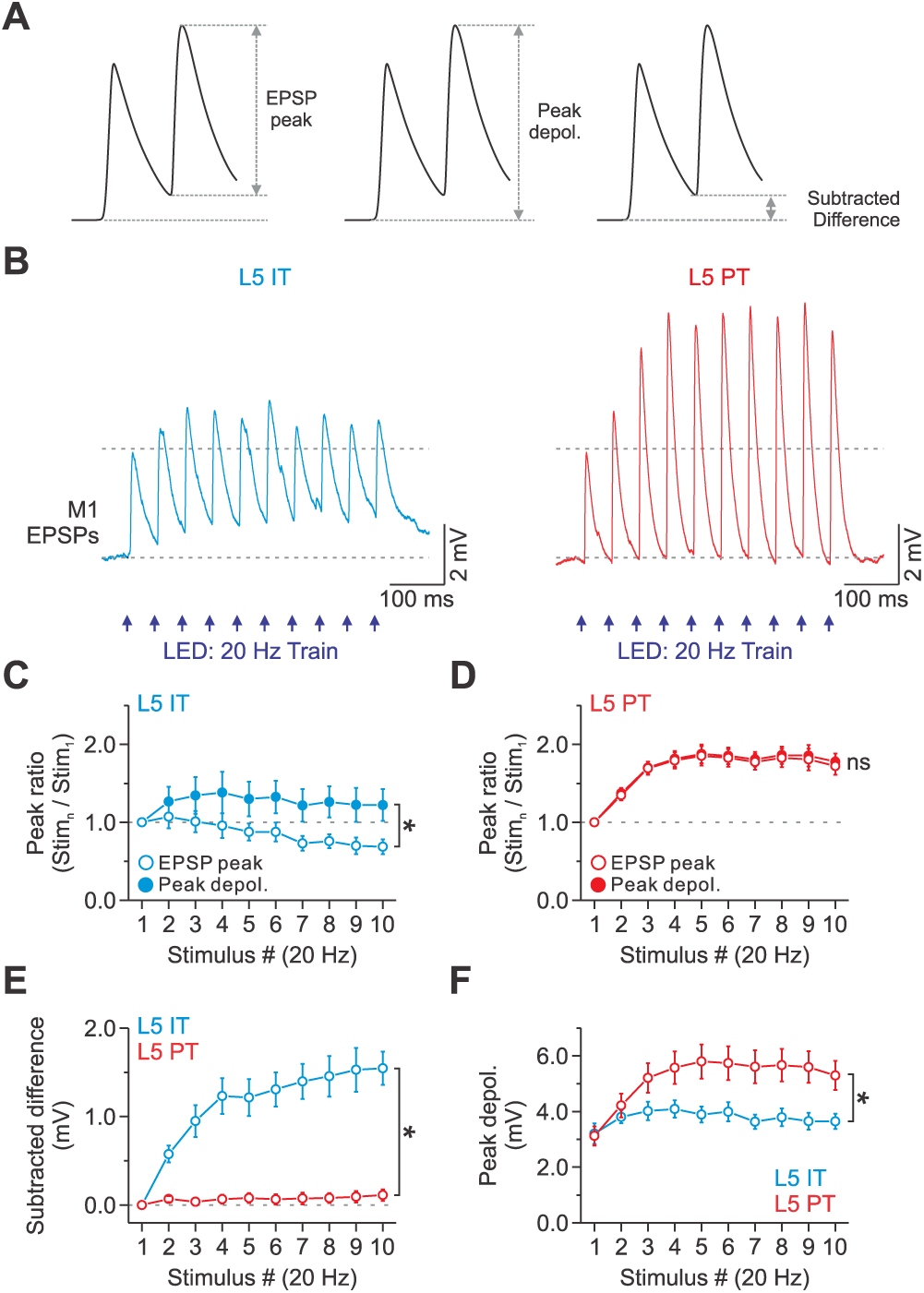
Trains of M1 input excite L5 PT neurons more strongly than IT cells due to short-term facilitation. **A**, The panel shows representative M1 responses and the methods used to calculate the EPSP peak, peak depolarization, and subtracted difference. **B**, Average EPSPs evoked by a 20 Hz train of optical stimuli recorded in a single L5 IT (Left) and PT neuron (Right). M1 responses were recorded at -94 mV in current-clamp, near the reversal for inhibition, and the light intensity was set to obtain an initial sub-threshold EPSP of ∼3 mV. **C**, Summary plot showing the average EPSP peak and peak depolarization ratio as a function of stimulus number within trains (normalized to the first response) for IT neurons. Note the EPSP peaks decreased during repetitive stimulation, whereas the peak depolarization increased through the train (p = 1.79E-6, two-way ANOVA, stim. 2–10, n = 5 cells from 3 mice). **D**, Summary plot showing the average EPSP peak and peak depolarization ratio as a function of stimulus number within trains (normalized to the first response) for PT neurons. There is no significant difference between the EPSP peak and the peak depolarization (p = 0.54533, two-way ANOVA, stim. 2–10, n = 24 cells from 11 mice). **E**, Summary plot showing how much the subtracted difference accounts for the peak depolarization as a function of stimulus number within trains for IT and PT populations (p = 3.91E-64, two-way ANOVA, stim. 2-10, n = 5 IT cells from 3 mice and 24 PT cells from 11 mice). **F**, Summary plot showing the average peak depolarization in millivolts as a function of stimulus number within trains for IT and PT populations (p = 2.08E-4, two-way ANOVA, stim. 2–10, n = 5 IT cells from 3 mice and 24 PT cells from 11 mice). Values are represented as mean ± SEM.

In contrast, because M1-evoked EPSPs decay more quickly in PT neurons, we found no overlap of consecutive EPSPs, resulting in no difference between the EPSP peak and the peak depolarization (**Figure 5D**). Thus, the depolarization associated with the preceding M1-evoked EPSP (the subtracted difference between the peak depolarization and EPSP peak) does not contribute to the increasing response amplitude in PT neurons but does in IT neurons (**Figure 5E**). In other words, temporal summation of EPSPs was minimal in PT neurons but significant in IT neurons. However, despite minimal summation in PT neurons, the M1 pathway was much more effective at exciting L5 PT neurons than IT cells during a short 20 Hz train (**Figure 5F**). This is because the elicited responses in PT neurons approximately doubled across the 20-Hz train due to short-term facilitation, whereas the responses in IT cells increased by only about 35% due to a combination of summation and short-term depression. Across the examined population, PT neurons had approximately 50% larger M1-evoked responses than the IT cells by the fifth pulse in the train.

Thus, M1 inputs elicited distinct postsynaptic responses in L5 PT and IT neurons that differ in time course, short-term dynamics, and how they integrate. These data also indicate that short-term synaptic facilitation significantly enhances M1 response amplitudes in PT neurons during repetitive 20 Hz stimulation with minimal contribution due to temporal summation.

Given the strength of M1 responses increase in PT neurons during repetitive activation, we next performed separate current-clamp experiments to determine how facilitating input impacts AP firing in these neurons. Excitatory M1 synapses target not only the basal dendrites of L5 pyramidal neurons but also their apical tuft in L1 (Petreanu et al., 2009), where they can interact with backpropagating action potentials (bAPs) to trigger large amplitude calcium spikes, leading to high-frequency burst firing in PT neurons (Stuart and Sakmann, 1994; Schiller et al., 1995; Larkum et al., 1999a; Larkum et al., 1999b; Grewe et al., 2010). As such, short trains of M1 synaptic input might couple more effectively with bAPs to trigger bursts of APs in the soma of PT neurons. To test this, we next performed current-clamp recordings from the somata of labeled PT neurons to examine how optical M1 stimuli couple with somatic action potentials. We chose to test POMp neurons as our PT population because these cells showed the highest propensity to burst when depolarized with positive current injections (**Table 1**). All cells were injected with direct current to produce a similar resting membrane potential at the soma (approx. -77 mV), and synaptic inhibition was intact throughout the recording. When M1 axons were activated in isolation using a single, low-intensity light pulse (LED_s_), somatic responses were always subthreshold (2.2 ± 0.3 mV, n = 10 cells; **Figure 6A**). Short 20 Hz trains (LED_T_) of the same light intensity were also subthreshold, rarely generating somatic APs (only 1 of 10 cells or 1 of 61 total trials; **Figure 6B**), whereas brief positive current injections at the soma (I_soma_; 1-35 ms, typically 5 ms; 0.35-1.85 nA) reliably evoke single APs (61 of 61 total trials; **Figure 6C**). For PT neurons, LED_T_ of M1 synaptic input coupled with a somatic AP could elicit a high-frequency burst of 3-4 APs (3.3 ± 0.3 APs; frequency: 213.4 ± 13.8 Hz) at the soma of some PT neurons (4 of 10 cells or 23 of 61 total trials), resulting in a significantly greater mean spike output than pairing a somatic AP and a single M1 synaptic input (I_soma_: 1.0 ± 0.0 APs; I_soma_+LED_S_: 1.1 ± 0.1 APs; I_soma_+LED_T_: 2.0 ± 0.3 APs; **Figure 6D-F**). For most PT neurons, coupling an LED_s_ (same intensity) and a somatic action potential was ineffective in evoking a high-frequency burst (observed bursting in only 1 of 10 cells or 1 of 61 total trials).

**Figure 6.**
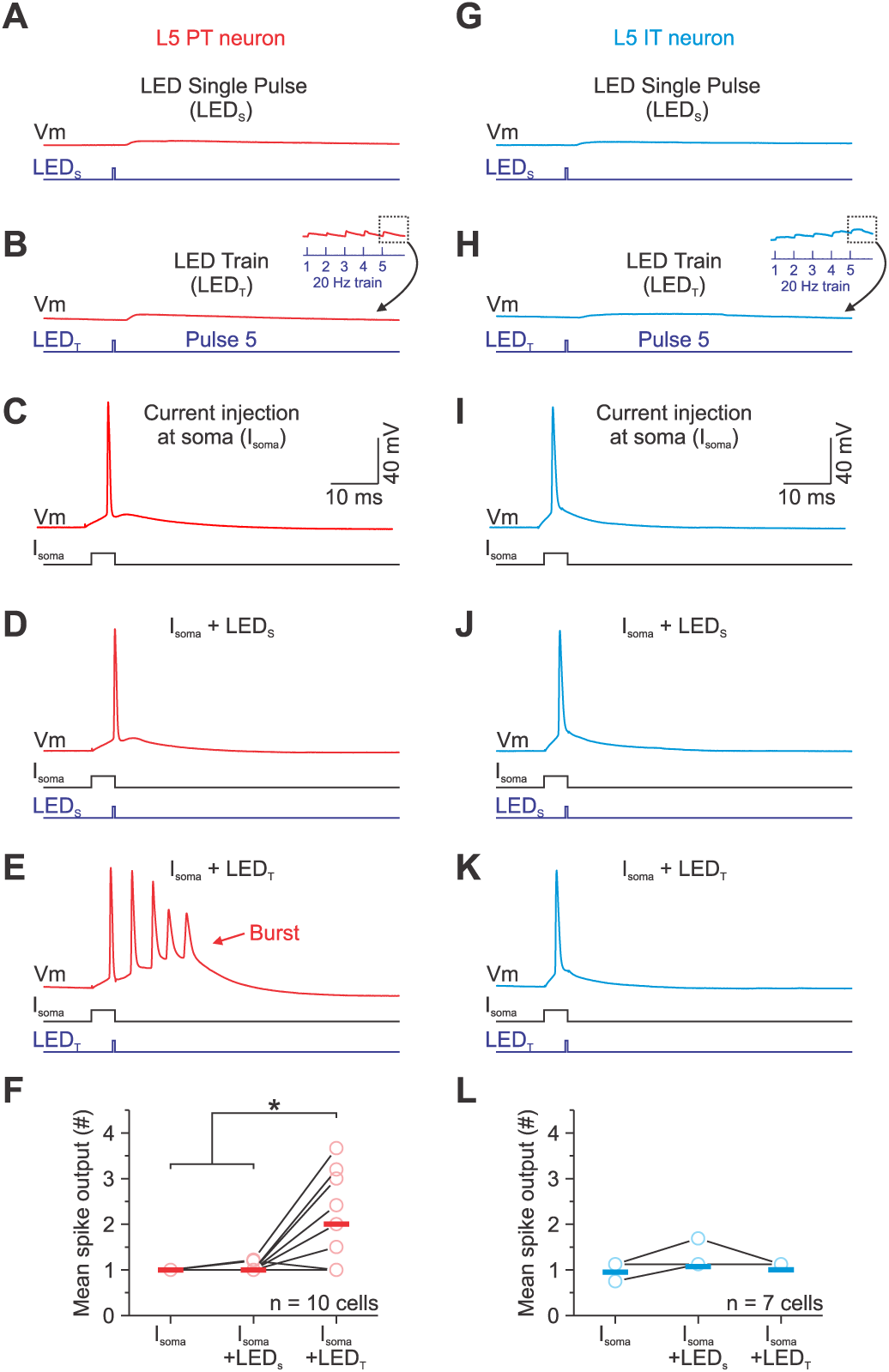
Coupling a short train of M1 synaptic inputs with a single spike at the soma increases the spike output and burst probability in L5 PT neurons. **A-B**, Subthreshold EPSPs evoked by a single LED stimulus (**A:** LED_s_) or the 5^th^ pulse of a 20 Hz train of stimuli (**B:** LED_T_) for an example L5 PT neuron. The LED_s_ and the first response in an LED_T_ only produced a voltage response of 2-3 mV at the soma and never reached the threshold for either an action potential or calcium-mediate action potential. (**C**) A short threshold current injection (typically 5 ms) at the soma (I_soma_) evoked a single action potential. (**D-E**) Voltage responses to combining somatic action potential (used in **C**) with a single EPSP (used in **A**) or the EPSP evoked by the 5^th^ pulse of a 20 Hz train (used in **B**) separated by an interval of 3-4 ms between the start of the somatic current injection and that of the light pulse. The mean synaptic latency from the onset of the light was 2.3 ± 0.1 ms (n = 10 cells; 7 mice). **F**, Group data summarizing the action potential output at the soma during coupling for L5 PT neurons. Coupling a somatic action potential with the 5^th^ pulse of an optical train produced significantly more spikes (* indicates p < 0.006, one-way ANOVA). **G-L**, Same as **A-F** for an example L5 IT neuron. There was no difference in spike output at the soma during coupling for L5 IT neurons (p = 0.258, one-way ANOVA) or the number of cells bursting for the M1p cells. The bars in **F** and **L** represent the means.

Unlike PT cells, IT neurons cannot generate burst firing at the soma (Moberg and Takahashi, 2022). However, L5 IT cells can generate bAPs, although their propagation is less efficient than PT cells (Grewe et al., 2010). Thus, we repeated the same experiments in a subset of IT cells to determine if bAPs in these neurons interact with M1 synaptic inputs to change somatic output. In contrast to L5 PT cells, combining an LED_s_ or LED_T_ with a somatic AP did not significantly change their overall spike output for L5 IT neurons (I_soma_: 1.0 ± 0.0 APs; I_soma_+LED_S_: 1.1 ± 0.1 APs; I_soma_+LED_T_: 1.0 ± 0.0 APs, n = 7 cells, **Figure 6G-L**).

Previous work has shown that initiating a burst of APs with a single distal input is highly time-dependent, requiring dendritic input and APs to coincide within 3-7 milliseconds (Larkum et al., 1999b). Given the time course of M1-evoked EPSPs did not change through a 20 Hz train, we hypothesized that EPSPs late in a train would still need to coincide with APs within a narrow time window to generate a burst at the soma. To test this, we delivered an LED_T_ of M1 synaptic input at different intervals after a single I_soma_. Consistent with our hypothesis, we found that the optimal time for generating a burst with an LED_T_ was very narrow, typically 0-4 ms after the AP (or 2-6 ms when accounting for a 2 ms synaptic latency; **Figure 7**). For intervals of 10 ms or greater, the probability of eliciting a burst with an LED_T_ fell quickly (**Figure 7B-C**). Altogether, these results demonstrate that M1 input can facilitate bursts in PT neurons when coupled with bAPs, that sustained M1 activity can increase the probability of this coupling due to strong short-term facilitation of M1 synapses, and that repeated M1 activation does not influence the timing-dependence of this coupling due to the fast time course of individual M1-evoked EPSPs.

**Figure 7.**
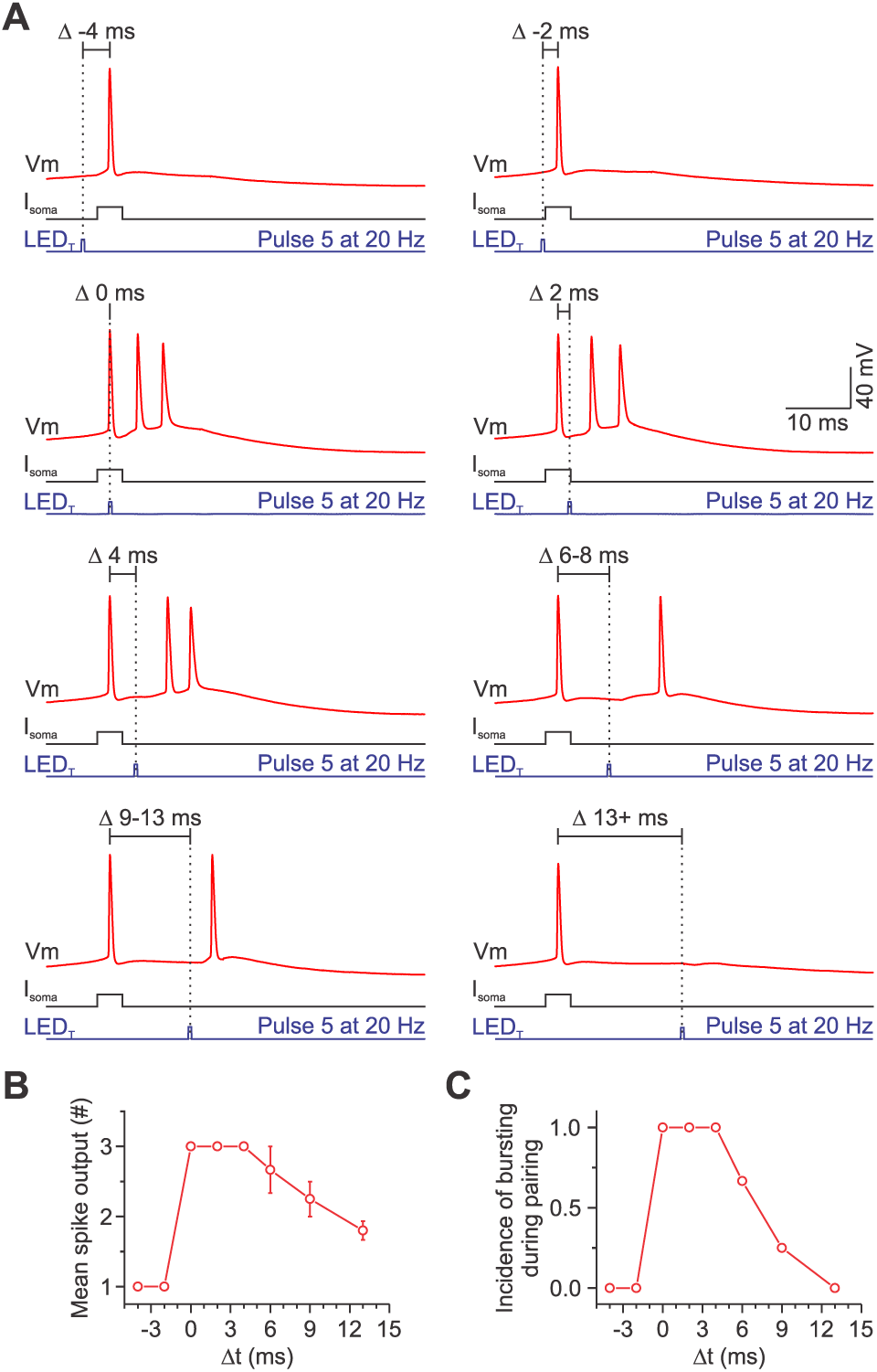
Coupling a train of M1 synaptic inputs with a somatic action potential requires precise timing in PT neurons. **A** , Voltage responses of a POMp neuron to combining a somatic action potential with the EPSP evoked by the 5^th^ pulse of a 20 Hz train separated by different time intervals (-4, -2, 0, 2, 4, 6-8, 9-13, and 13+ ms). The actual timing was based on the peak of the AP and not the onset of the current injection. Optically evoked synapse responses had synaptic delays with short onset latencies (∼2 ms). **B-C**, Plots show that this cell produced more spikes and had a higher incidence of bursting when the onset of the 5^th^ pulse was 0-6 ms after the somatic action potential.

## DISCUSSION

One of the most prominent structural features of the neocortex is the presence of numerous corticocortical pathways interconnecting areas (Rockland and Pandya, 1979; Felleman and Van Essen, 1991). While it is generally thought that these pathways convey contextual information, such as attention, expectation, and action command (Gilbert and Li, 2013), we are only beginning to understand how these inputs influence their target regions. Realizing the nature of these influences requires information about the physiological properties of the connections themselves and how specific cell types respond to those signals. Here, we targeted the synapses and specific cells involved in M1-S1 interactions, revealing that M1 connections to distinct classes of L5 neurons in S1 are dramatically different.

Although the net connection strength of M1 was similar between classes of L5 neurons, during repetitive activation, we found that excitatory M1 synapses onto IT cells displayed modest short-term depression, whereas M1 synapses onto PT cells showed robust short-term facilitation. We also found that the response to single M1 stimuli had a slower time course in IT neurons than PT cells, partly due to differences in Ih expression. Despite the short-term depression observed in IT neurons, the slower kinetics of M1-evoked EPSPs resulted in greater temporal summation and a postsynaptic potential that steadily increased during short trains. In contrast, we found minimal temporal summation in PT neurons due to the faster time course of M1-evoked EPSPs. However, because the short-term facilitation of M1-evoked EPSPs was strong in PT neurons, M1 responses continued to increase for subsequent stimuli. Although the overall postsynaptic potential increased in both cell types during repetitive M1 activation, M1 responses were ∼50% stronger in PT neurons than IT cells by the fifth stimulus in trains. Thus, the consequences of the reduced temporal summation in PT neurons are outweighed by the synaptic facilitation of the excitatory M1 synapse itself. Functionally, because EPSPs within trains of M1 stimulation were strong and fast in PT neurons, we found that they coupled more effectively with bAPs within a narrow time window to generate somatic bursts in these cells. Thus, we conclude that there are two parallel but dynamically distinct systems of M1 synaptic excitation in L5 of S1, each defined by the short-term plasticity of its chemical synapses, the projection class of the postsynaptic target neuron, and how the postsynaptic neuron responds to those inputs. These physiological differences in the connections between M1 and S1 L5 excitatory circuits may be specializations related to their particular functions during sensorimotor processing.

Selective optogenetic stimulation of axons has been used to study postsynaptic targets, synapse locations, strength, and the impact of M1 input on various excitatory and inhibitory cells in S1 (Petreanu et al., 2009; Lee et al., 2013; Zagha et al., 2013; Kinnischtzke et al., 2014; Kinnischtzke et al., 2016; Shen et al., 2022). However, our understanding of their short-term dynamics has lagged due to the complications with using opsins to study synapses (Jackman et al., 2014). Recently, our lab has addressed these challenges and found that M1-evoked responses in S1 facilitated with repetitive activity. While this is true for most M1 to S1 connections, we found responses in a subset of L5 neurons that displayed short-term depression (Martinetti et al., 2022).

The present study is the first to demonstrate that this diversity in short-term dynamics depends on the postsynaptic L5 neuron and their axonal projection target, with M1 synapses onto IT cells depressing and those onto PT cells facilitating. The relevance of such variable short-term dynamics in corticocortical projections is unclear. At synapses where depression is common, there is a decrease in synaptic strength with activity (Zucker and Regehr, 2002). Previous work has suggested that depressing intracortical synapses provide a dynamic gain-control mechanism that enables them to respond proportionally to the percent change in firing frequency rather than the absolute change (Abbott et al., 1997). Thus, during periods of elevated M1 activity, such as bouts of whisking, depressing signals may enable L5 IT cells to encode relative changes in M1 firing. In future studies, targeting other IT neurons, such as claustrum or callosal projecting cells, would be interesting to determine if they have similar synaptic properties. In contrast, since facilitating synapses are usually weak initially but enhance synaptic strength with activity (Jackman and Regehr, 2017), PT cells may be less responsive to transient M1 stimuli but ideally suited to respond to sustained M1 activity or a greater number of synchronous M1 inputs.

Importantly, M1 inputs onto IT neurons in other layers facilitated with repetitive activity, indicating that short-term depression is a unique property of synapses connecting with L5 IT neurons and not a general rule of M1 connections to all IT cells in S1. These findings provide physiological evidence for cell-type and layer-specific corticocortical connection specificity at a synaptic physiology level. These results complement previous findings of target-cell-specific short-term dynamics in local cortical circuits (Markram et al., 1998; Reyes et al., 1998; Beierlein et al., 2003; Lefort and Petersen, 2017) and highlight how the specificity and synaptic dynamics of connections also govern long-range neuronal circuits underlying the interactions between cortical areas.

The diversity in short-term dynamics found here is consistent with earlier studies investigating the pathways linking secondary and primary sensory cortices and the callosal pathway within the prefrontal cortex (Covic and Sherman, 2011; De Pasquale and Sherman, 2011; Lee et al., 2014). However, two of these studies (Covic and Sherman, 2011; De Pasquale and Sherman, 2011) suggest that corticofugal neurons receive depressing inputs, contrasting the present results. We also found that the net synaptic strength of M1 input was similar across L5 neurons, consistent with previous studies examining the pathways linking M1 to S1 and the retrosplenial cortex to the secondary motor cortex (Kinnischtzke et al., 2016; Yamawaki et al., 2016). Conversely, in visual cortices, long-range corticocortical inputs preferentially target looped L5 IT neurons (Young et al., 2021). Whether these discrepancies are due to methodological differences or represent distinct connectivity features between these pathways is unclear. Indeed, recent studies using similar optical strategies have begun revealing key differences between corticocortical pathways (Naskar et al., 2021; Martinetti et al., 2022; Shen et al., 2022). Thus, further studies are needed to determine if corticocortical synapses onto L5 neurons follow general or specialized rules depending on the area and projection type.

In addition to receiving inputs with distinct temporal dynamics, we also found L5 neurons respond to M1 signals differently. Specifically, the M1-evoked EPSP time course was faster in PT neurons than IT cells, with Ih partially responsible for these cell-type-specific differences. Functionally, the slower kinetics increased the summation during short trains of M1-evoked EPSPs in IT cells, whereas the faster time course reduced the summation in PT neurons. These results are consistent with work highlighting the importance of Ih in influencing the kinetics and propagation of synaptic responses in L5 pyramidal neurons (Magee, 2000; Williams and Stuart, 2000; Berger et al., 2001; Harnett et al., 2015) and the idea that PT neurons express higher levels of Ih compared to IT cells (Dembrow et al., 2010; Sheets et al., 2011; Dembrow et al., 2015; Anastasiades et al., 2018). Of course, we cannot exclude the possibility that M1-mediated feedforward inhibition also influenced the time course of EPSPs (Pouille and Scanziani, 2001; Gabernet et al., 2005; Mittmann et al., 2005). Furthermore, we cannot rule out additional physiological properties, such as differences in whole-cell capacitance, differential synapse locations, expression of synaptic receptors, or other voltage-activated ion channels (Kumar and Huguenard, 2003; Petreanu et al., 2009; Lafourcade et al., 2022). Nevertheless, the differences in M1-evoked EPSP time course imply that IT neurons behave more as temporal integrators, integrating temporally dispersed M1 synaptic events over a broad time window, whereas PT neurons behave as coincidence detectors, even over sustained periods of activity, summating temporally coincident inputs over a narrow time window (Konig et al., 1996). Functionally, it has been proposed that neurons operating as temporal integrators and coincidence detectors are well suited to perform rate and synchrony coding, respectively (Ratte et al., 2013; Dembrow et al., 2015).

M1 axons projecting to S1 terminate densely in L1, where they make connections with the apical tuft dendrites of L5 neurons (Cauller et al., 1998; Veinante and Deschenes, 2003; Petreanu et al., 2009). Although single, local depolarizing events in the dendrites undergo significant voltage attenuation (Rall, 1967; Williams and Stuart, 2002), distal dendritic depolarization coupled with single bAPs can trigger dendritic calcium spikes and somatic bursts (Larkum et al., 1999b). Such somatodendritic coupling has been proposed as a cellular mechanism through which L5 neurons implement translaminar input associations (Larkum, 2013). Consistent with this hypothesis and recent work (Shen et al., 2022), we showed that M1 synaptic inputs can trigger bursting in PT neurons when paired with temporally precise somatic spikes. Moreover, we demonstrate that the probability of generating somatic bursts increased with sustained M1 input due to presynaptic facilitation while maintaining a narrow time window for somatodendritic coupling, most likely due to fast EPSP kinetics. The short-term dynamics of M1 synapses onto PT neurons suggest that they are ideally suited for robust transmission over relatively extended periods, consistent with the sort of activity observed during voluntary whisking. In contrast, we found that coupling bAPs with either low- or high-frequency M1 input did not change the spiking output of IT cells.

The results we describe here indicate that the M1 to S1 corticocortical projection comprises two parallel but dynamically distinct systems of synaptic excitation in L5, each defined by the short-term plasticity of its synapses, the postsynaptic cell type targeted, and the integration of its inputs. Because of their unique dynamic properties, distinct L5 projection neurons are likely to be differentially engaged by patterns of M1 activity that vary in frequency and timing. Our work complements previous *in vivo* work demonstrating the importance of motor-associated synaptic inputs and active dendritic processing in generating L5 activity as well as the differences in activity between IT and PT neurons during active whisker-mediated behavior (Xu et al., 2012; Manita et al., 2015; Ranganathan et al., 2018; Takahashi et al., 2020; de Kock et al., 2021).

## Acknowledgments

We thank Dr. Charles Lee Cox and the Michigan State University Center for Advanced Microscopy for imaging support and Luis E. Martinetti for stereotactic virus injection support.

## Author Contributions

H.H.K and S.R.C designed research; H.H.K, K.E.B, and G.G performed research; D.M.A, T.K., and H.H.K performed the histology and reconstructions. H.H.K and S.R.C analyzed the data; H.H.K and S.R.C wrote the paper.

---

**Extended Data Figure 1.**
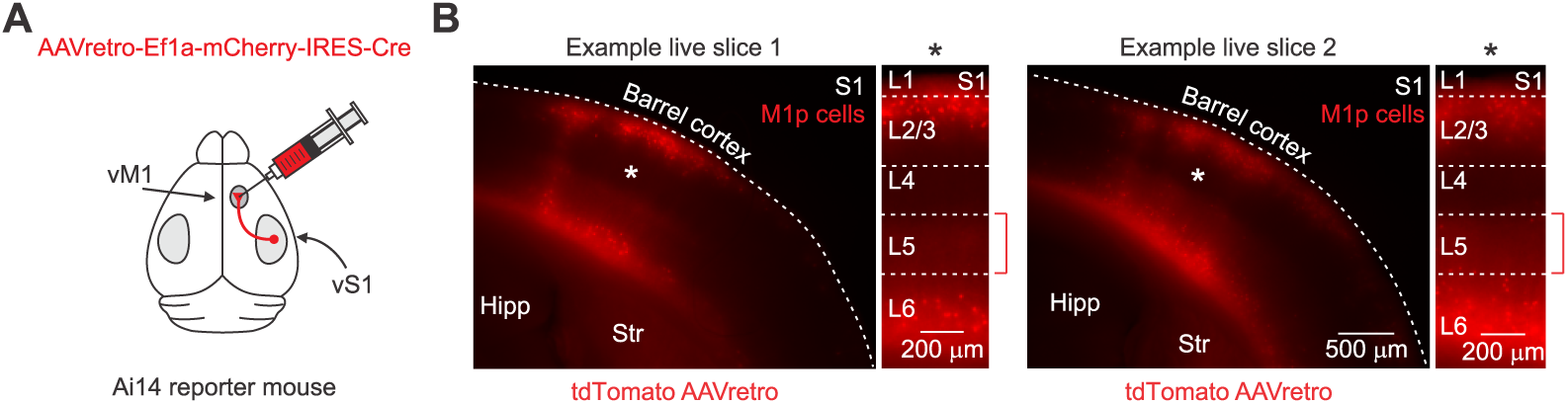
Anatomical characterization of L5 IT neurons labeled with AAVretro. **A**, Injection schematic showing AAVretro carrying genes for Cre and mCherry was injected unilaterally into M1 of Ai14 mice *in vivo* at ∼3 weeks of age. **B**, Two example fluorescent images of live coronal slices (300 μm thick) through S1 from two different Ai14 mice injected in M1 ∼21 days prior with AAVretro.EF1a-mCherry-IRES-Cre. Images show tdTomato expressing S1 neurons following AAVretro injection. Higher magnification images show retrogradely labeled neurons in L2/3 and L6, but very few in L5 (n = 5 mice).

